# Novel modifications to fructose metabolism are associated with adaptation to high-sugar diets in bats

**DOI:** 10.64898/2026.09.29.755168

**Authors:** J.M. Jamison, A.M. Gutierez, P. Oelbaum, G. Marron, A. Sadier, J. Ortega, F. Reid, N. Simmons, K.C. Welch

## Abstract

High-sugar diets in mammals are associated with a number of metabolic diseases including diabetes and obesity. However, frugivorous and nectivorous bats specialize in consuming diets high in simple sugars while remaining healthy. To investigate the metabolic adaptations in bats with high-sugar diets, we conducted targeted metabolomics on blood samples collected during the first ten minutes after glucose or fructose feeding in five bat species whose natural diets vary in sugar content. By comparing changes in blood metabolite levels over time both between feeding treatments and across dietary guild, we have been able to identify shifts in sugar processing that likely support the healthy metabolism of simple sugars mammals with high-sugar diets. Notably, despite both sugars being readily absorbed, only fructose feeding induced significant changes in other metabolites within ten minutes after feeding. Species who naturally consume more sugar showed both faster production of metabolic intermediates and lowered production of common deleterious byproducts of fructose metabolism than species with comparatively lower-sugar diets. The prioritization and refinement of fructose metabolism is therefore likely a more key aspect of how bats with high-sugar diets are able to avoid the diseases typically associated with excessive sugar consumption than modifications to glucose metabolism.

## Introduction

Carbohydrates are commonly found in mammalian diets in a variety of forms such as insect exoskeletons, plant components, and even within vertebrate muscle [1]. However, these carbohydrates typically exist as polymers which must be broken down into monomeric sugars before they can be absorbed into the body [2,3]. This delay of digestion and competition for absorption with other nutrients slows the absorption of monosaccharides [4]. Consuming large quantities of glucose in its monomeric form can cause rapid increases in glucose concentrations in the blood and cells, where high concentrations of the sugar can cause cellular dysfunction through the non-enzymatic glycation of proteins. Liver and muscle cells, which absorb can sequester glucose into glycogen, where its glycative potential is limited. Some other cells can use the polyol pathway to turn glucose into sorbitol and subsequently fructose, but the enzymes of this pathway are not expressed in all cell types. Furthermore, sorbitol creation causes osmotic stress and cell bursting, and the generation of fructose depletes NADPH in the cell (which is necessary to regenerate the antioxidant glutathione) while creating a sugar fourteen times more glycative than the original glucose (Figure 1).

**Figure 1.**
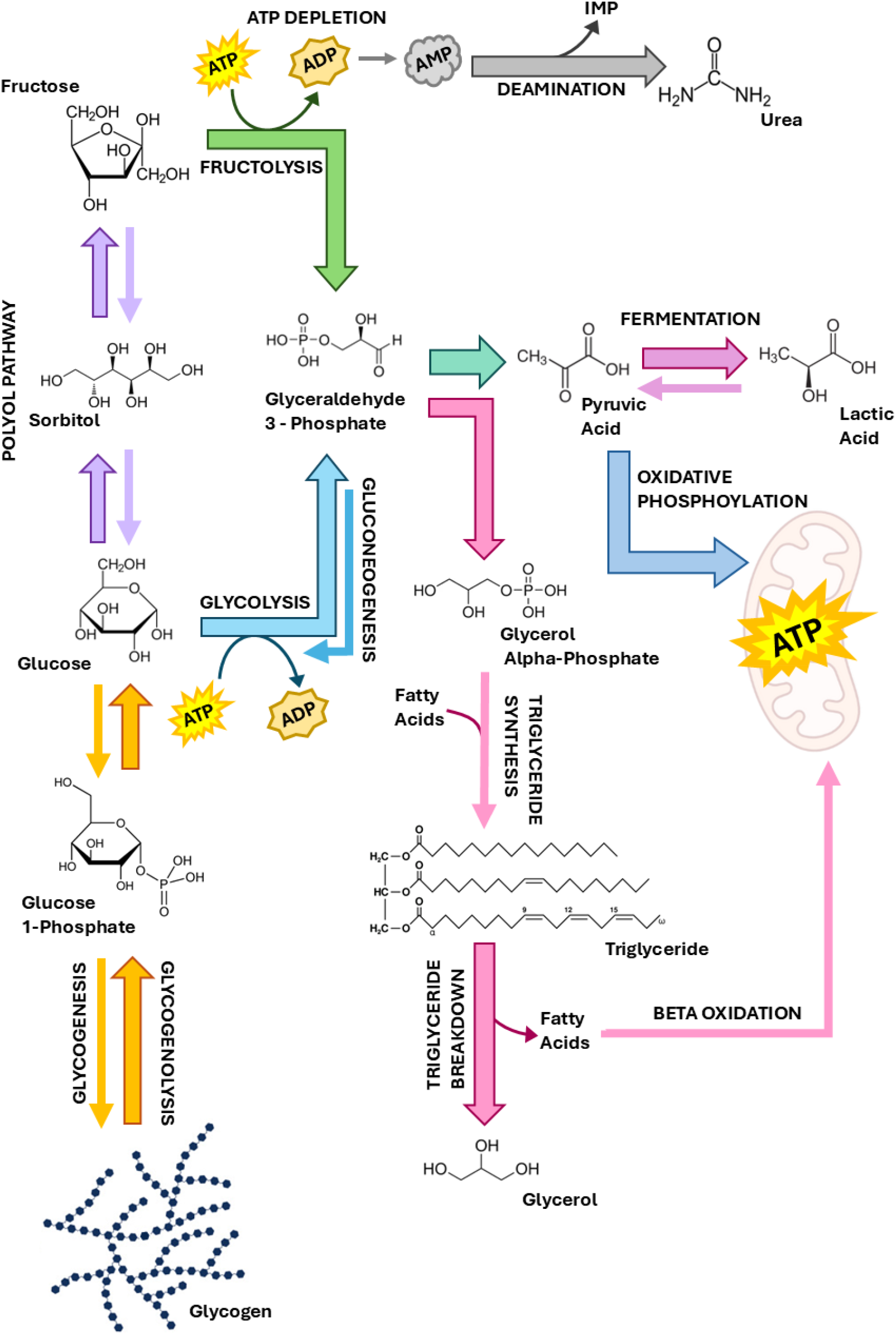
Simplified metabolic pathway interactions of measured metabolites. Pathways are color coded and identified in all capital letters. Primary pathways of investigation are shown with block arrows while secondary associated pathways are shown with arrow lines. Names of metabolites are indicated in sentence case next to the respective molecular structure. Oxidative phosphorylation and beta oxidation occur within the mitochondria [15], but are labelled on arrows pointing to the mitochondria.

This process, in addition to hormone dysregulation, is a key part of why diets high in monomeric glucose, such as the Western diet, are associated with metabolic diseases. The Western diet additionally contains levels of fructose never seen before in human evolution [5]. In low doses, nearly all fructose consumed is converted into triose-phosphates in the intestinal epithelia which are used for energy within the cell. In high doses, however, this intestinal filter gets overwhelmed, and fructose enters the blood stream where it begins wreaking havoc in the body [6]. Fructose is a uniquely glycative sugar, with high-fructose diets resulting in significantly more whole-body non-enzymatic glycation even compared to diets high in glucose or sucrose [7]. Breaking down fructose may remove its glycative harm, but metabolizing large quantities of fructose results in ATP depletion and uric acid accumulation (which is associated with cardiovascular disease and gout), because fructolysis is a completely unregulated pathway that uses ATP stores faster than oxidative phosphorylation can replenish it (Figure 1). Large amounts of fructolysis additionally lead to fatty deposits in the liver (where almost all fructolysis occurs) as one of its intermediates, fructose-1-phosphate, promotes de novo lipogenesis, making excess energy be stored preferentially as fat, not glycogen [8]. This explains why high-fructose, but not high-glucose diets, have been shown to enhance lipid deposition in the liver, contributing to non-alcoholic fatty liver disease (NAFLD), which in turn is a precursor to hepatic steatosis, metabolic syndrome, and type II diabetes [9–11].

While the metabolic processes that result in high levels of simple sugar consumption leading to disease in humans are well known, the changes in metabolism that nectivorous and frugivorous bats have evolved that allow them to healthily process large quantities of simple sugars every day remains poorly understood. Several species of these bats have been shown to maintain blood glucose levels over eight times higher than those of a healthy human after feeding [12]. But the mechanisms by which these animals are able to function with blood glucose concentrations that would put a human in a coma are still unknown. Molecular evolution analyses have identified dozens of genes related to the transport and metabolism of glucose and fructose that are under positive selection in nectivorous bats, but the specific effects of the identified mutations on metabolic flux are still a mystery [13].

To elucidate adaptations associated with high-sugar diets in bats, we performed targeted metabolomics on bat plasma before and at five or ten minutes after the feeding of either a glucose or fructose meal. Bats rapidly utilize exogenous sugars to fuel their metabolism, making the minutes immediately after feeding of particular interest to characterize this unique metabolic system [14]. To generate a comprehensive picture of how glucose and fructose are metabolized in bats, we measured the flux of ingested sugar (glucose or fructose), the amount of energy generation (via lactic and pyruvic acid levels), the breakdown of endogenous fuel stores (via glucose-1-phosphate, glycerol, and glycerol-alpha-phosphate levels), and the production of products associated with the polyol pathway and fructolysis (sorbitol and uric acid respectively). We compared these values in bat species adapted to diets along a gradient of sugar incorporation, from insectivory to specialized nectivory.

## Methods

### Ethics Statement

All experiments were carried out under University of Toronto Animal Ethics Protocol #20012113. Samples were collected at Lamanai Archeological Reserve, Orange Walk, Belize in April 2023 and April 2024 with approval from the Belize Forestry Department (FD/WL/1/23 (01); FD/WL/1/24(06)) and Institute of Archaeology (IA/H/1/23(01); IA/H/1/23 (02); IA/H/1/24(01); IA/H/1/24(02)). Additional field work was performed at Sylvan Camp and Falls, Puntenaeras, Costa Rica in March 2023 and March 2024 with permission of the Comisión Nacional para la Gestión de la Biodiversidad (R-014-2022-OT-CONAGEBIO). Work done at Tamana Cave, Trinidad in April 2023 was authorized by the Ministry of Agriculture and Fisheries. Samples collected in Mexico in August 2024 at Rancho Cacachilas, Baja California Sur, Mexico was done with permission of the Secretariat of Environment and Natural Resources (SGPA/DGVS/03632/2).

### Bat Capture

Animals used in Belize and Costa Rica were captured using a combination of ground and triple high mist nets (6m, 9m, 12m, 18m). Nets were opened shortly before dusk in front of flowers (often banana flowers) and along trails to maximize bat capture during foraging. We checked nets for captured animals at intervals of 5-10 minutes from the time the nets opened until they were closed at approximately 23:00. Any bats that were trapped by the nets were immediately removed and placed into cotton bags until the commencement of experiments. Pregnant individuals were immediately released.

For cave roosts sampled in Mexico and Trinidad, we entered the cave shortly before dusk, while the animals were active but before they had exited for the evening. Using a hand net, we captured bats flying inside of the cave. Upon capture, individuals were identified to the species level, and non-pregnant individuals of the target species were placed individually into cotton bags until the beginning of experiments. All other individuals of non-target species were immediately released.

### Blood Sampling

All individuals were fasted for at least one hour after capture before sampling commenced. Due to the small size of the animals, only one blood sample was taken per individual. Males and females were split as equally as possible between glucose and fructose treatments and among timepoints (Table S1). Each blood sample was taken either before feeding, 5 minutes after feeding, or 10 minutes after feeding, with the meal fed consisting of a 20% w/w aqueous solution of either glucose or fructose of a volume that equated the bat consuming 5.4 g sugar/kg bodyweight [14]. To collect blood samples for bats under 30 g, we cleaned the elbow of the individual with an ethanol wipe and allowed the ethanol to evaporate. Then, we used a 27 g heparinized needle to make a small puncture wound in the brachial vein in the bend of the elbow. For bats over 30 g, we first cleaned the top of the wing with an ethanol wipe and waited for the ethanol to evaporate. We then used a 25 g heparinized needle to pierce the cephalic vein on the underside of the wing, being careful not to pierce through to the dorsal side [16].

For both procedures, we used a heparinized micropipette tip to remove the blood as it beaded from the wound and transfer it to a heparinized microcentrifuge tube. We collected blood until the wound stopped bleeding or 150 μL of blood was collected for bats over 30 g or 70 μL of blood was collected for bats under 30 g, whichever happened first. In instances where the collection limit was reached first, we stopped the bleeding by applying pressure with a clean cotton pad to encourage clotting.

After collection, we spun the blood for 2 minutes at 6,000 rpm with a Sprout® mini centrifuge (HeathrowScientific^©^, USA) at ambient temperature to separate plasma from the red blood cells. We then transferred the plasma to a clean microcentrifuge tube. Both plasma and red blood cells were then flash frozen in liquid nitrogen. At the completion of the procedure, the bat was given ad libitum sucrose solution in the case of frugivorous and nectivorous bats and ad libitum water in the case of insectivorous bats. Bats were returned to their bags to rest and were released before dawn near their point of capture. All bats flew away readily at the end of the experiments.

### Metabolomics

We sent plasma samples to the West Coast Metabolomics Core at the University of California Davis for metabolomics analysis. There they underwent gas chromatography time of flight (GC-TOF) mass spectrometry on a Leco Pegasus IV mass spectrometer connected to a Restek© Rtx-5Sil MS column that was 30 m long with a 0.25 mm internal diameter. The column was filled with 0.25 μm film made of 95% dimethyl/5% diphenylpolysiloxane and guarded by a 10 m long empty guard column which was cut at 20 cm intervals if the reference quality control mixture indicated possible column contamination. For chromatography, 0.5 μL of each plasma sample was injected into the column via a multi-baffled glass liner at 50 °C which was then increased to 250 °C at a rate of 12 °C/s. The temperature of the oven was held at 50 °C for one minute then increased to 330 °C at a rate of 20 °C/min and then held at 330 °C for 5 minutes. The column utilized a helium mobile phase at a rate of 1 mL/min, which gives narrow peak widths of 2-3 s for primary metabolite classes. The unit mass resolution of the spectrometer was set to 17 spectra/s from 80-500 Da at -70eV ionization energy. The detector voltage was 1800 V with a 230 °C transfer line and a 250 °C ion source.

The spectra were preprocessed using ChromaTOF v2.32 without smoothing to deconvolute mass spectra at a signal/noise level of 5:1, resulting in 3 second peak widths with baseline noise subtracted off. A BinBase filtering algorithm (rtx5) was then used to determine apex masses. This algorithm removed both spectra containing less than 10 peaks with intensities greater than 10^7^ counts/second, indicating low quality and spectra with MS similarity greater than 800, which served to cap the intensity range for high m/z marker ions. The retention index was calculated by 5^th^ order polynomial regression. The spectra were cut to 5% base peak abundance and matched to database entries starting with the most abundant spectra and moving to the least. To be considered valid, spectra were additionally required to have a retention index window of ± 2000 units (about ± 2 seconds). Any unique ions that were identified had to be included in the reference index of masses and exist in more than 3% of base peak abundances to be considered. Finally, the mass spectrum similarity had to fit criteria dependent on peak purity, signal/noise ratios, and a final isomer filter. For each metabolite, peak heights were converted to concentrations based on a metabolite-specific nine-point calibration curve of known-concentration serial dilutions.

### Statistical Analyses

For each of the nine targeted metabolites, we verified that the starting concentrations were not significantly different between the glucose and fructose fed groups by performing a two-way ANOVA with the aov function in the stats v.3.6.2 package in R v 4.5.0 looking at the effect of species and sugar group overall and the effect of sugar group within each species on the metabolite concentration at timepoint zero [17]. To determine which species pairs differed significantly in baseline concentration for each metabolite, we performed pairwise t-tests between the species’ concentrations at timepoint zero and applied a Holm correction using the pairwise.t.test function from the stats R package.

To evaluate the effect of feeding glucose or fructose on metabolite levels over time, we then conducted a two-way ANOVA using the aov function in the stats package testing the effects of species and timepoint across the whole dataset and the effect of species within each timepoint on the metabolite concentrations [17]. For metabolites where species and/or timepoint were significant we conducted a pairwise t-test using the pairwise.t.test function in the stats R package which grouped the metabolite concentrations by each significant parameter. In cases where the interaction of time and species was significant, we performed a pairwise t-test looking at the difference in species’ means independently for each timepoint. For each metabolite where significant impacts of species, time, and/or their interaction were found, we collated all p-values from the t-tests done for that metabolite and applied a false discovery rate correction for multiple testing using the p.adjust function in the stats R package.

For t-tests that indicated a significant effect of species across all timepoints or at a single timepoint, we used a simulation-based approach to verify if the difference between species was more extreme than that which would be expected based purely on phylogeny. For each significant comparison, we used the data to calculate the average rate of Brownian motion (σ^2^) across the tree using the pic function from the ape v5.8-1 R package [18]. We then used this σ^2^ value to simulate data on the tree 480638 independent times using ape’s fastBM function. The function simulates only one value per species, so we clustered the simulations together in groups of six to mimic the sample size of the empirical data. We then repeated the significant t-tests using the simulated data for the species being compared for every group of 6 simulations. To find the probability that our observed data was more extreme than the simulated data, we totaled the number of t-tests where the simulated t-value was larger than the observed t-value and divided that by the number of t-tests run. To correct for the multiple tests being run on the data, we corrected the final p-values with a Holm correction that adjusted the family-wise type I error for the number of p-values calculated for that metabolite.

We additionally conducted ANOVAs that looked at the difference in levels of metabolites between glucose and fructose feeding for each species. To do this, we used the aov function from the stats R package to test the impact of sugar fed and timepoint overall and the impact of sugar fed at each timepoint on each metabolite. Because there were only two fed conditions, we did not need to run subsequent t-tests when a metabolite was significantly affected by the type of sugar fed. Finally, we performed pairwise t-tests with the pairwise.t.test function in the stats R package to test the differences in each metabolite concentration between timepoints for each species [17]. We corrected the p-values with a Holm adjustment independently for each test.

### Phylogenetic Tree Creation

To correct statistical analyses for the effect of phylogeny, we first generated a phylogenetic tree from Cytochrome B (CYTB) gene sequences using the topology of (Table S2)[19]. For Glossophaga mutica, we used the sequence of Glossophaga soricina that was collected most closely to the field site in Belize, as G. soricina was recently split into several species, and the database names have not been updated [20]. We similarly placed G. mutica in the location of G. soricina in the Rojas (2016) topology. To generate branch lengths for the tree, we aligned the CTYB sequences with MUSCLE v3 in SeaView v5.0.5 and input this alignment with the tree topology to baseml in paml v4.9 [21–23]. The resulting tree was used in all phylogenetic corrections (Figure S1).

## Results

### Metabolite Responses to Sugar Consumption

When bats were fed glucose, significant changes occurred in plasma concentrations of glucose, fructose, sorbitol, and glucose-1-phosphate across all species while the levels of the other measured metabolites did not significantly change from their pre-feeding values (Table S6). Plasma glucose values increased the most rapidly and dramatically of the metabolites in all species, increasing on average from 1422.56 ± 234.06 ng/μL to 6653.90 ± 293.64 ng/μL by five minutes after feeding (Table 1). *Macrotus californicus* had the least extreme change in plasma glucose, but this species had significantly higher starting plasma glucose levels than the other species, which is typical of mammals who consume primarily proteins (Figure 2A) [24]. Considering only the species who regularly consume predominantly carbohydrates, plasma glucose concentrations increased on average by 700 ng/μL in just five minutes after feeding a glucose solution (Table 1). In most species, plasma glucose concentrations either decreased or remained the same between five and ten minutes after feeding, though the decrease was only significant in *Glossophaga mutica* (p = 0.027). However, even with this general lowering, the plasma glucose values remain extrordinarily high across all species. This includes one *Phyllostomus discolor* individual who reached a plasma glucose concentration of 10,284 ng/μL, the highest value in the dataset, though several other individuals of this species exceed plasma glucose levels of 10,000 ng/μL.

**Table 1.** Average metabolite values (ng/μL) with standard deviations for each species either before feeding or at five or ten minutes after glucose feeding.

| Species | Time (min) | Glucose | Fructose | Lactic Acid | Pyruvic Acid | Sorbitol | Urea | Glucose-1-Phosphate | Glycerol-Alpha-Phosphate | Glycerol |
| --- | --- | --- | --- | --- | --- | --- | --- | --- | --- | --- |
| *Anoura geoffroyi* | 0 | 840 ± 321 | 10 ± 7 | 876 ± 293 | 95 ± 51 | 2 ± 1 | 192 ± 190 | 1,478 ± 2,239 | 20 ± 4 | 74 ± 39 |
| | 5 | 5,889 ± 2,441 | 49 ± 49 | 448 ± 123 | 123 ± 73 | 18 ± 8 | 341 ± 168 | 371 ± 241 | 20 ± 10 | 67 ± 31 |
| | 10 | 6,496 ± 1,235 | 33 ± 18 | 227 ± 101 | 101 ± 41 | 18 ± 8 | 426 ± 130 | 232 ± 145 | 21 ± 4 | 75 ± 21 |
| *Glossophaga mutica* | 0 | 776 ± 886 | 10 ± 4 | 736 ± 570 | 114 ± 68 | 3 ± 2 | 364 ± 180 | 813 ± 622 | 18 ± 4 | 80 ± 20 |
| | 5 | 7,703 ± 1,011 | 29 ± 13 | 714 ± 249 | 105 ± 63 | 24 ± 4 | 492 ± 226 | 1,096 ± 1,527 | 18 ± 5 | 71 ± 31 |
| | 10 | 6,443 ± 774 | 26 ± 11 | 956 ± 743 | 171 ± 177 | 22 ± 9 | 408 ± 205 | 388 ± 289 | 20 ± 7 | 115 ± 88 |
| *Carollia castanea* | 0 | 925 ± 639 | 5 ± 1 | 372 ± 147 | 71 ± 45 | 6 ± 4 | 527 ± 258 | 11,370 ± 9,897 | 11 ± 2 | 57 ± 26 |
| | 5 | 6,703 ± 825 | 27 ± 22 | 345 ± 197 | 97 ± 22 | 17 ± 12 | 619 ± 272 | 870 ± 665 | 13 ± 4 | 35 ± 15 |
| | 10 | 5,182 ± 1,989 | 12 ± 5 | 368 ± 145 | 90 ± 51 | 12 ± 7 | 548 ± 319 | 2,379 ± 4,226 | 12 ± 5 | 22 ± 11 |
| *Phyllostomus discolor* | 0 | 1,115 ± 787 | 7 ± 5 | 811 ± 398 | 104 ± 19 | 2 ± 1 | 310 ± 198 | 4,675 ± 5,772 | 13 ± 2 | 21 ± 10 |
| | 5 | 7,329 ± 1,472 | 29 ± 16 | 756 ± 494 | 111 ± 63 | 12 ± 5 | 306 ± 181 | 939 ± 430 | 20 ± 8 | 11 ± 6 |
| | 10 | 6,909 ± 2317 | 35 ± 28 | 766 ± 264 | 99 ± 41 | 11 ± 5 | 315 ± 126 | 810 ± 677 | 16 ± 5 | 12 ± 11 |
| *Macrotus californicus* | 0 | 3,507 ± 1,265 | 41 ± 25 | 849 ± 574 | 119 ± 67 | 3 ± 1 | 1,220 ± 161 | 123 ± 78 | 16 ± 6 | 71 ± 24 |
| | 5 | 5,819 ± 1,091 | 35 ± 15 | 775 ± 287 | 183 ± 83 | 5 ± 2 | 1,182 ± 342 | 130 ± 78 | 18 ± 4 | 49 ± 16 |
| | 10 | 6,883 ± 868 | 43 ± 11 | 539 ± 313 | 110 ± 70 | 7 ± 2 | 1,174 ± 249 | 326 ± 282 | 13 ± 3 | 57 ± 31 |

**Figure 2.**
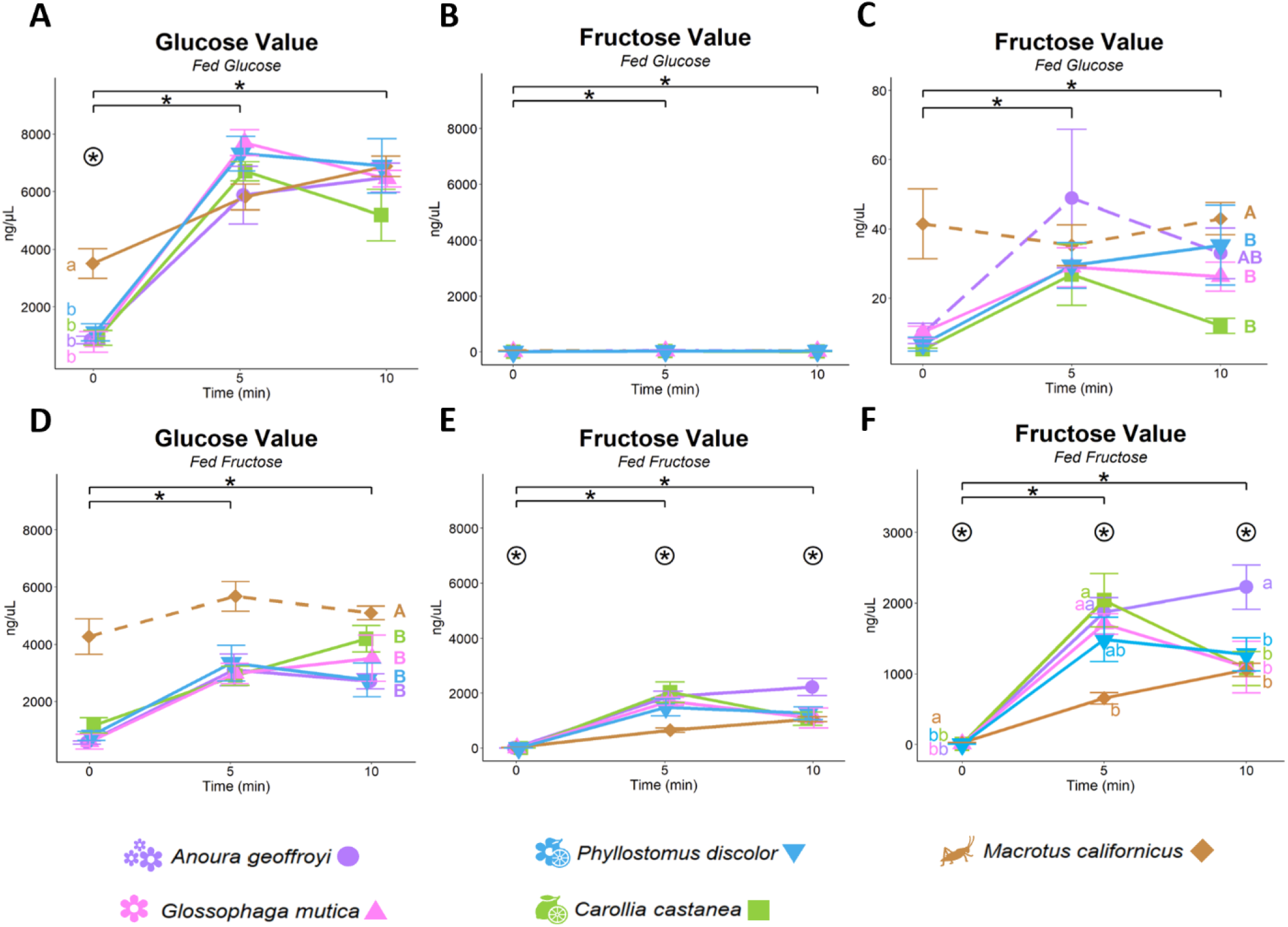
Changes in concentrations (ng/μL) of A) plasma glucose after glucose feeding, B&C) plasma fructose after glucose feeding, D) plasma glucose after fructose feeding, and E&F) plasma fructose after fructose feeding. Panels A,B, D and E are set to the same scale. Panels C&F are zoomed in version of panels B&E respectively to allow differences between species to be better shown. Asterisks with brackets indicate significant differences across all species between the indicated timepoints. Asterisks within circles indicate significant differences across species within a timepoint, as grouped by the lowercase letters. Line type indicates significant differences across all timepoints between species as grouped by capital letters. n = 6 individuals per sugar per species per timepoints.

Surprisingly, plasma fructose values also increased significantly after glucose feeding across all species, though by a far less extent than plasma glucose, though this was driven primarily by the changes in the non-insectivorous species, as *M. californicus* showed no significant differences in plasma fructose concentration between any of the timepoints (Figure 2C, Table S8). None of the tested species experienced significant changes in the inidcators of energy production, pyruvic acid and lactic acid, within ten minutes of consuming the glucose meal (Figure 3). However, after feeding there was significant decline in glucose-1-phosphate levels, particularly in C. castenea and P. discolor, suggesting a decrease in glycogenolysis (Figure 4A).

**Figure 3.**
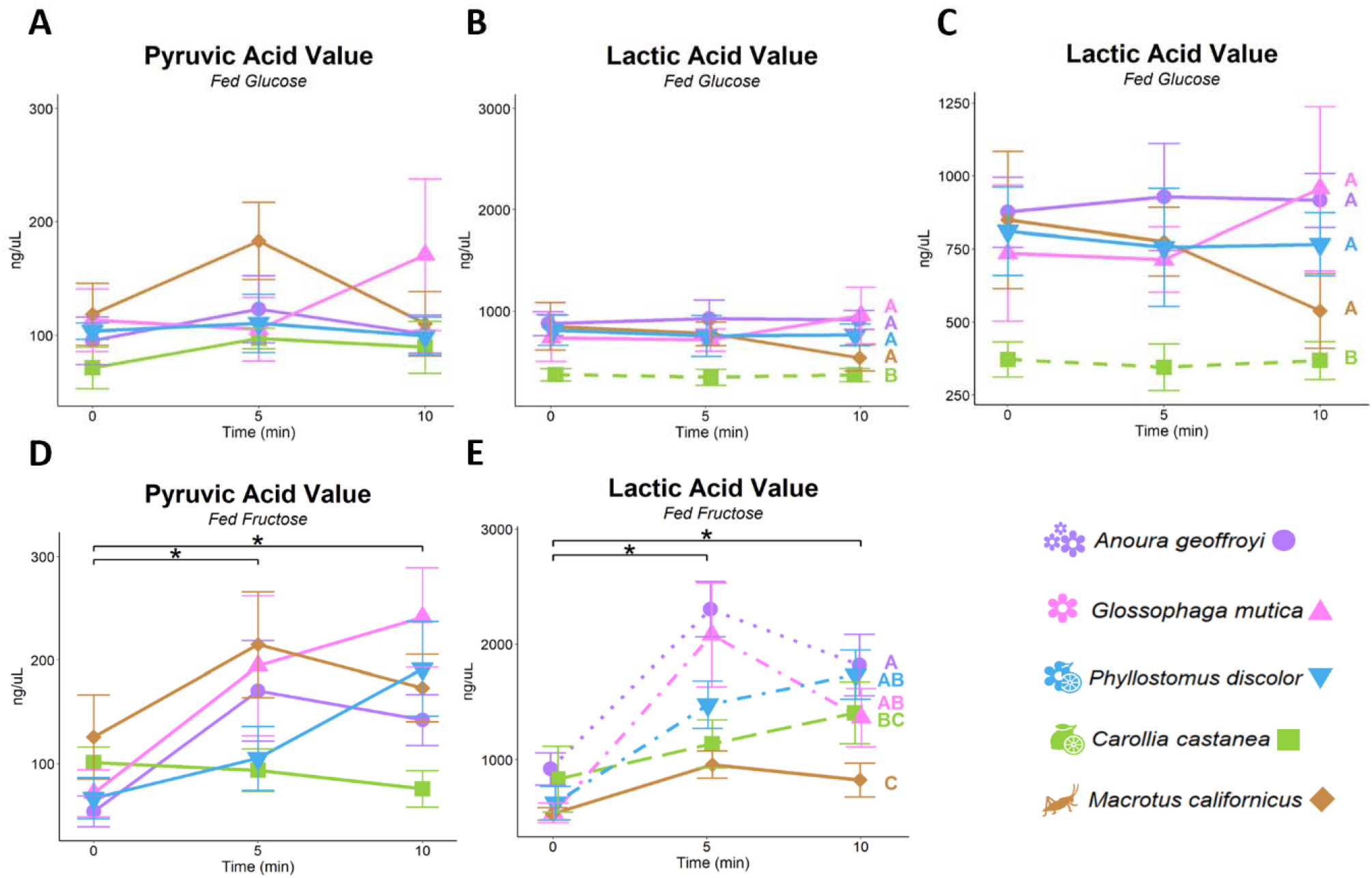
Changes in plasma concentrations (ng/μL) of A) pyruvic acid after glucose feeding, B&C) lactic acid after fructose feeding, D) pyruvic acid after fructose feeding, and E) lactic acid after fructose feeding. Panels A&D and B&E are set to the same scale. Panel C is a zoomed in version of panel B allowing differences between species to b more easily visualized. Asterisks with brackets indicate significant differences across all species between the indicated timepoints. Line type indicates significant differences across all timepoints among species as grouped by capital letters. n = 6 individuals per sugar per species per timepoint.

**Figure 4.**
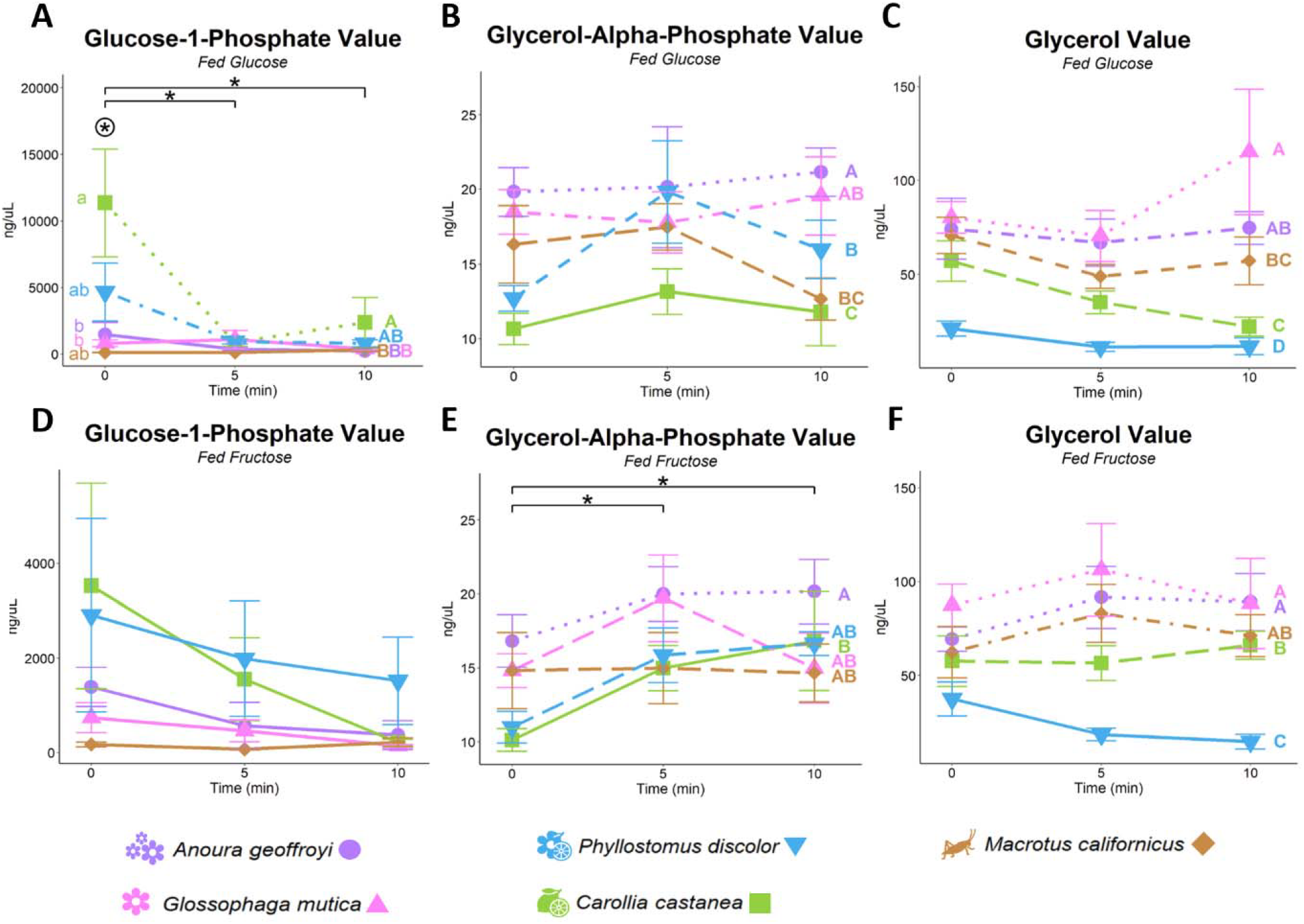
Changes in plasma concentrations (ng/μL) of A) glycerol after being glucose feeding, B) glycerol after fructose feeding, C) glycerol-alpha-phosphate after glucose feeding, D) glycerol-alpha-phosphate after fructose feeding, E) glucose-1-phosphate after glucose feeding, and D) glucose-1-phosphae after fructose feeding. Panels A&B and C&D are set to the same scale. Asterisks with brackets indicate significant differences across all species between the indicated timepoints. Asterisks within circles indicate significant differences across species within a timepoint, as indicated by the lowercase letters. Capital letters indicate significantly different values averaged across all timepoints among species. n = 6 individuals per sugar per species per timepoint.

Plasma urea levels also showed no significant changes with glucose feeding, but *Macrotus californicus* consistently had significantly higher concentrations of urea in its plasma than any other species across all timepoints, with the average difference between it and the other species being 792.51 ± 63.88 ng/μL (Figure 5A). Sorbitol concentrations did increase significantly from baseline values across species after the consumption of glucose (Figure 5B&C). At five minutes post-feeding, the two nectivorous species, *Glossophaga mutica* and *Anoura geoffroyi* had the largest increases in sorbitol concentration, followed by the frugivorous *Carollia castanea* and the semi-nectivorous *Phyllostomus discolor*, and finally the insectivorous *Macrotus californicus*. At five minutes only the differences between the nectivorous and insectivorous species were significant, and at ten minutes the only differences that were significant were those between *A. geoffroyi* and *P. discolor* and *A. geoffroyi* and *M. californicus. M. californicus* showed by far the smallest increase in plasma sorbitol after glucose feeding of all the species with the average change between baseline levels and levels at timepoint ten being 3.83 ± 1.01 ng/μL compared to the pooled average for the other species at 13.00 ± 1.77 ng/μL.

**Figure 5.**
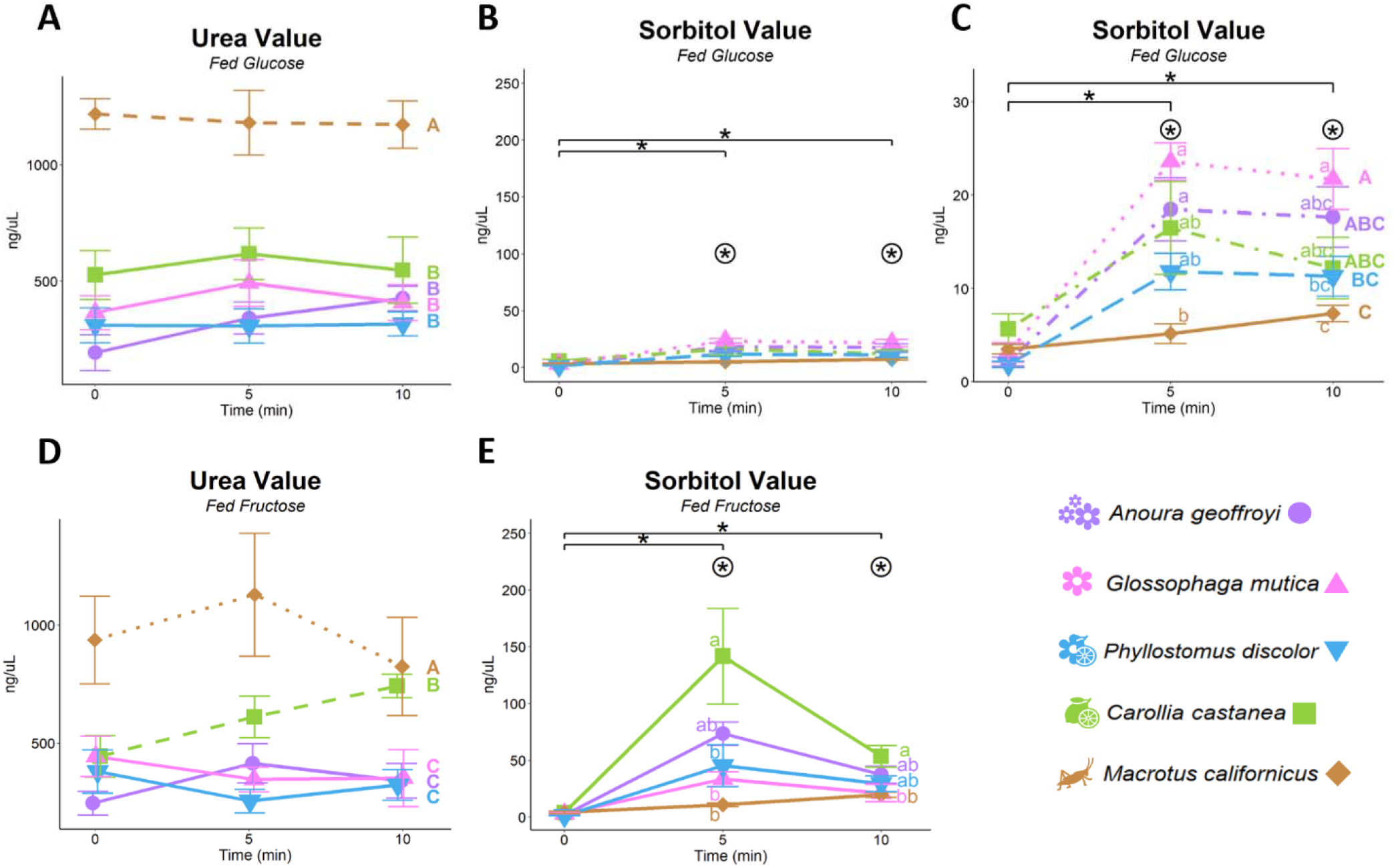
Changes in plasma concentrations (ng/μL) of A) urea after glucose feeding, B&C) sorbitol after glucose feeding, D) urea after feeding fructose, and D) sorbilol avter fructose feeding. Panels A&D and B&E are set to the same scale. Panel E is a zoomed in version of panel C allowing differences between species to be more easily visualized. Asterisks with brackets indicate significant differences across all species between the indicated timepoints. Circled asterisks indicate timepoints where significant differences exist between species, as grouped by the lowercase letters. Line types indicate significant differences across all timepoints between species, as grouped by the capital letters. n = 6 individuals per sugar per species per timepoint.

Fructose feeding resulted in changes in more of the measured metabolites in the first ten minutes after consumption than glucose feeding, with only urea and glucose-1-phosphate concentrations not significantly changing over time across species (Table S17). A greater number of metabolites also showed significant differences among species after fructose feeding compared to glucose consumption (Table 2). Glycerol was the only metabolite that did not have a significant or marginal p-value after testing for the impact of species on its concentration over time after fructose feeding (Table S17).

**Table 2.**
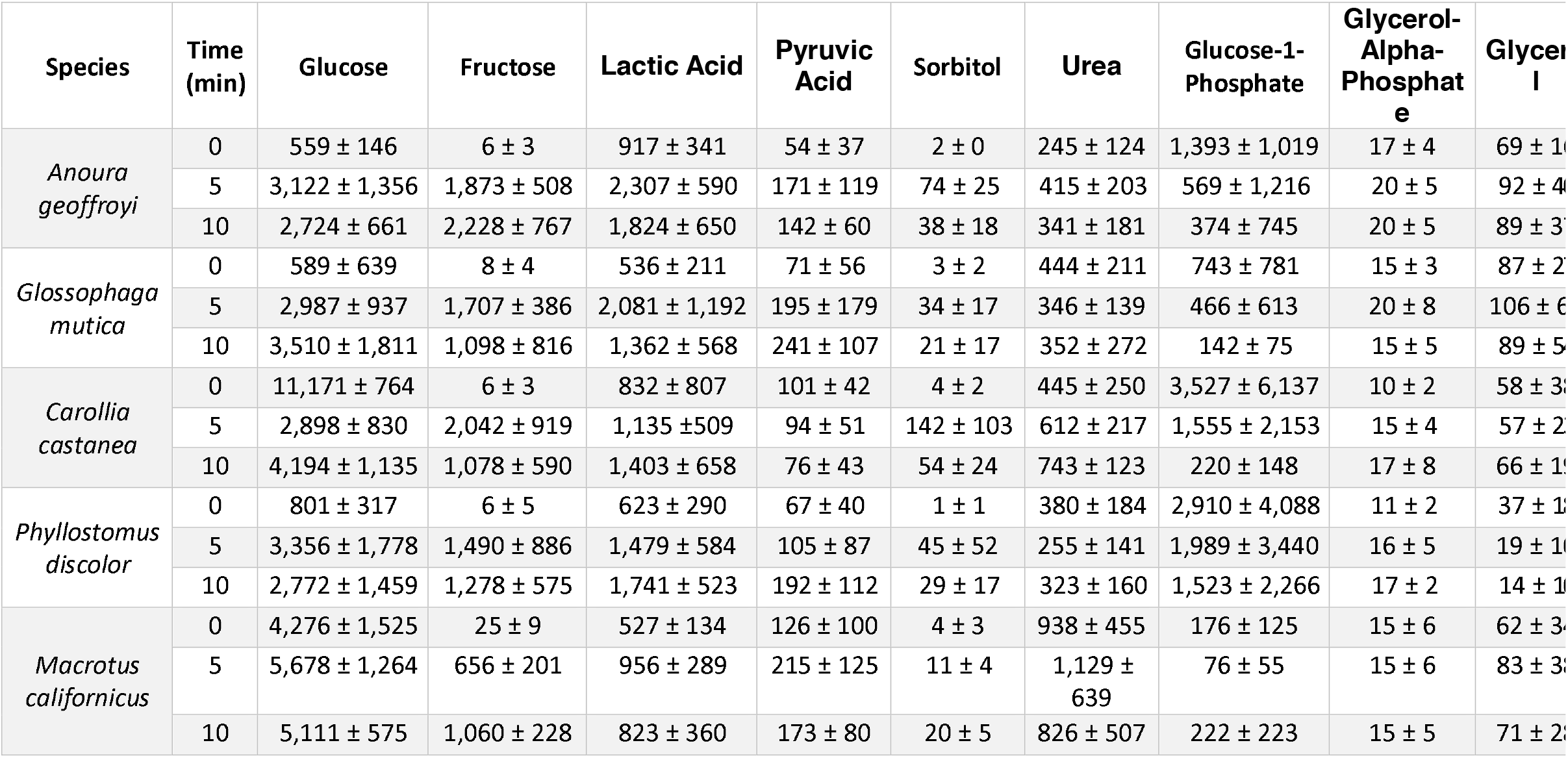
Average metabolite values (ng/μL) with standard deviations for each species either before feeding or at five or ten minutes after fructose feeding.

The most striking difference in the metabolite profiles across species after fructose feeding when compared to glucose feeding is the difference in plasma fructose concentrations. All species had significantly higher plasma fructose concentrations after consuming fructose than they did after eating glucose (Table S17). By five minutes after feeding, the non-insectivorous species reached plasma fructose concentrations averaging 1,754.07 ± 136.35 ng/μL, nearly 52 times higher than the plasma fructose concentration five minutes after glucose feeding and almost 265 times higher than baseline blood fructose levels. Even the insectivorous *Macrotus* californicus showed significant increases in blood fructose levels. By ten minutes after feeding, *M. californicus* had plasma fructose concentrations that were not significantly different from most of the other species at the ten-minute timepoint (Figure 2F). The exception is the specialized nectivore Anoura geoffroyi whose blood fructose levels continued to rise between five to ten minutes, unlike the other non-insectivorous bats that showed decreases in or maintained similar plasma fructose concentrations. Due to the continued increase in plasma fructose levels, at ten minutes after fructose feeding, *A. geoffroyi* reached plasma fructose levels double most of the other species (2,227.83 ± 313.03 ng/μL). Plasma glucose also rose significantly in the ten minutes after fructose feeding, despite no glucose being ingested. This was primarily driven by the non-insectivores, though this was a significantly smaller rise than these species experienced after glucose feeding (Table S19). The plasma glucose levels of the non-insectivores were significantly lower than those of *M. californicus* across all timepoints, despite *M. californicus*’ plasma glucose levels not significantly changing from its baseline values (Table S20).

Unlike with glucose feeding, both indicators of sugar breakdown, pyruvic acid and lactic acid, increased significantly over time across species after fructose feeding (Table S17). While the overall increase in pyruvic acid pooled across species did significantly increase over time, none of the increases within species were significant, though Anoura geoffroyi had a marginal p-value (p = 0.069) for the difference in pyruvic acid concentration between five and ten minutes after feeding (Table S20). Both nectivorous species, *A. geoffroyi* and G. mutica, exhibited large increases in lactic acid at five minutes, which then decreased by the ten-minute timepoint, while P. discolor and C. castanea showed slower, continuous increases in lactic acid levels over time (Figure 3E). *M. californicus* showed minimal overall changes in lactic acid concentration (Table 2).

Levels of urea in the bats’ plasma did not significantly change over time after fructose feeding across all species (Table S18). However, *C. castanea* clearly showed a substantial increase in plasma urea concentrations after fructose feeding that was nearly statistically significant (p = 0.053). This increase was sufficient for *C. castanea* to have a significantly higher average value for plasma urea across timepoints compared to other non-insectivorous bats, and by ten minutes after fructose feeding, *C. castanea’s* plasma urea levels are indistinguishable from those of the insectivorous *M. californicus* (Figure 5D). Therefore, *C. castanea* is likely unique in its processing of fructose when compared to the other species, which leads to this progressive increase in urea in the plasma. *C. castanea* was also unique in that it exhibited significantly higher levels of plasma sorbitol than almost every other species (except *A. geoffroyi*) at five minutes after fructose feeding (Table S6). Though all the species had significantly higher plasma sorbitol concentrations after fructose ingestion when compared to glucose ingestion except for *G. mutica* (Table S17).

Like with the glucose-fed group, individuals of P. discolor and C. castanea had the highest starting levels of glucose-1-phosphate, and these levels dropped over the course of the experiment, though these values were not significantly different from the other tested species potentially due to the large amount of inter-individual variation (Figure 4D). After fructose feeding, P. discolor and C. castanea showed p-values approaching statistical significance (p = 0.12 and p = 0.077 respectively) for the difference in the plasma concentrations of the lipogenesis indicator, glycerol-alpha-phosphate, between the start and end of the experiment (Table S20).

## Discussion

There are clear modifications to sugar metabolism in bats who are specialized on high-sugar diets compared to those who specialize on lower-sugar diets. Interestingly, these modifications appear to be more focused on the metabolism of fructose more so than glucose in the minutes immediately following sugar consumption. The absorption of glucose across all species was incredibly rapid, likely due to paracellular nutrient absorption, which is common among bats, with most species reaching maximum levels of plasma glucose at five minutes after feeding [25]. These levels are the highest ever recorded in mammals and exceed the upper range of most glucometers, emphasizing the importance of prioritizing early timepoints and diluting samples to accurately capture the scope of adaptation to high sugar diets in this group [12,26,27].

Despite the rapid rise in plasma glucose values after glucose feeding, these values remained substantially elevated for the duration of the experiment. Other studies of blood glucose after feeding in resting nectivorous bats indicate that blood glucose levels in some species remain elevated above 4,000 ng/μL for up to thirty minutes after glucose consumption [12,26]. This suggests that the movement of glucose into the tissues after intestinal absorption occurs at much slower time scale. Corroborating this is the failure of most of the metabolites we measured to show significant changes over time in any species following glucose feeding. There was a drop in circulating glucose-1-phosphate levels in P. discolor and C. castanea indicative of a reduction of glycogenolysis, but there was not a corresponding increase in lactic acid or pyruvic acid as one would expect if exogenous sugar was being metabolized for energy. The capabilities of these animals to withstand prolonged periods of hyperglycemia suggests a suite of adaptations that prevent the non-enzymatic glycation of plasma proteins. Such modifications have previously been identified in birds, which have naturally high blood sugar levels compared to mammals, but anti-glycative adaptations in plasma proteins have not been described in mammals to date.

While glucose is permitted to remain in the blood at high concentrations for prolonged periods of time, fructose is almost immediately either converted to glucose or broken down for energy after absorption in non-insectivorous bats. Blood glucose levels exceed blood fructose levels for all species except the specialized nectivore, *A. geoffroyi* (Figure 2). While these bats are tolerant to high blood glucose, it seems that they do not allow blood fructose to reach these same levels, perhaps because of the much more glycative nature of fructose when compared to glucose [28]. It is clear that the non-insectivorous species use the polyol pathway to convert fructose into sorbitol and likely glucose, based on the plasma sorbitol levels (Figure 4). This is the first record of the polyol pathway using fructose as a starting material in a mammalian system. The levels of sorbitol seen in *C. castanea* also suggest that the enzymes of the polyol pathway are far more widely expressed than they are in other mammals. While *C. castanea* has the highest sorbitol values, this cannot be easily interpreted as it using the polyol pathway the most. Sorbitol is difficult to move across cell membranes, resulting in osmotic stress and cellular bursting [29]. It is therefore likely that most of the sorbitol found in the blood is due to burst cells. The lower plasma sorbitol levels of the in the frugi-nectivorous and nectivorous species may be due to adaptations to better handle the osmotic stress induced by sorbitol creation rather than a lessened use of the polyol pathway overall. However, further investigation is needed to fully differentiate between these alternatives.

The movement of fructose into the cell for energy is clearly used by non-insectivorous bats to remove fructose from circulation, and the increases in pyruvic acid and especially lactic acid show that this is happening in non-insectivorous bats on a timescale of minutes after fructose feeding, unlike with glucose feeding (Figure 3). The breakdown of fructose occurs more rapidly in the nectivorous species than in the frugi-nectivorous or frugivorous species, suggesting that a more rapid breakdown of fructose is beneficial for diets higher in nectar, where large, sudden influxes of fructose are more common. Despite their rapid metabolism of fructose however, neither the nectivorous species nor the frugi-nectivorous species showed increases in the level of urea in the plasma, which would be expected with ATP depletion associated with unregulated fructolysis. These species, therefore, have either developed mechanisms to regulate fructolysis or to eliminate urea without exposing it to wider circulation, either of which would represent novel mechanisms of metabolism in mammals.

In addition to not showing the increase of urea associated with fructolysis and subsequent ATP depletion, the nectivorous species did not have increases in plasma glycerol-alpha-phosphate, which would be indicative of lipogenesis, the typical fate of fructose in mammals. *C. castanea* and *P. discolor* did show this sign of lipogenesis, suggesting that the nectivorous species have evolved to not convert ingested fructose into fat. This is despite the fact that these species appear to rely more on fat than glycogen as an endogenous energy store compared to the other non-insectivorous species, based on their relatively lower fasting levels of glucose-1-phosphate and higher fasting levels of glycerol and glycerol-alpha-phosphate (Figure 5). The failure of fructose feeding to induce lipogenesis in nectivorous species further suggests that these species have adapted to have regulatory mechanisms for fructolysis and potentially even modifications to how lipogenic transcription factors are impacted by fructolytic intermediates.

All of the non-insectivorous species show modifications to sugar metabolism compared to the insectivorous *M. californicus* indicative of adaptations related to their high sugar diets. Regarding glucose metabolism, these species show lower baseline plasma glucose levels and faster and higher rates of glucose absorption than *M. californicus*. However, most other metabolites did not show significant changes after glucose feeding, showing that at least immediately after ingestion, glucose metabolism is not prioritized by species adapted to consuming large quantities of this sugar. Rather, fructose is the simple sugar that these species rapidly convert into other metabolites after eating. As these sugars are typically consumed together in high-sugar diets, it is possible that the minimal use of glucose after eating evolved in non-insectivorous bats to allow fructose to supply most of the body’s energy needs and in doing so more rapidly remove the sugar from the body.

Fructose metabolism also shows an increasing number of modifications in species with increasing degrees of nectar in their diet, indicating that fructose metabolism is a primary pathway under selection during adaptations to high-sugar diets over glucose metabolism. This is perhaps not surprising with fructose’s recent identification as a “metabolic hazard” in mammals, making its metabolism a necessary detoxification for animals who consume large quantities of it [30]. The first adaptations evolved to aid in fructose detoxification include the use of the polyol pathway to rapidly convert fructose into glucose, as shown by increases in sorbitol being shared among all of the non-insectivorous bats. As nectar is incorporated into the diet, the polyol pathway is either relied on less to minimize osmotic stress or species evolve mechanisms to prevent cellular bursting due to the swelling induced by sorbitol creation. Similarly, the breakdown of fructose but not glucose immediately after ingestion aids in detoxifying fructose in non-insectivorous bats, with this process being even further accelerated once bats evolve nectivory (Figure 3E).

Having nectar as a significant part of the diet brings with it adaptations that regulate fructolysis or otherwise prevent urea resulting from ATP depletion from entering general circulation. It is only once true nectivory evolves, however, that the lipogenic effects of fructose become minimized. This is despite the fact that evolving nectivory also seems to coincide with a switch from these species depending primarily on glycogenolysis to get energy during a fast to lipolysis. Finally, the most specialized nectivores can evolve an increased tolerance for elevated blood fructose levels, with *A. geoffroyi* being the only species examined to not lower plasma fructose concentrations between five and ten minutes after feeding. While these conclusions need to be confirmed in other frugivorous and nectivorous species, this framework provides the first understanding of the molecular metabolic changes that accompany evolutionary increases in dietary sugar in mammals. It furthermore highlights the metabolic threat fructose poses to mammalian physiology while identifying the key modifications that allow some mammal species to consume large quantities of this sugar without developing metabolic diseases.

## Supporting information

Summplemental Figure

Supplemental Table Descriptions

## Acknowledgements

The authors wish to thank all of the local guides and field station staff that aided with in-countr logistics, particularly Pablo Guia and the staff at Rancho Cacahilas, Karen Gonzales and the Lamanai Outpost Team, Hammond Noreiga, and Francis Mora Zuñiga and Jesus Amado. We also want to acknowledge Scott Mackell, who oversaw the team at the UC Davis Metabolomics Core, which conducted the metabolomics analysis

