## Supplementary material for "Novel modifications to fructose metabolism are associated with adaptation to high-sugar diets in bats": Summplemental Figure

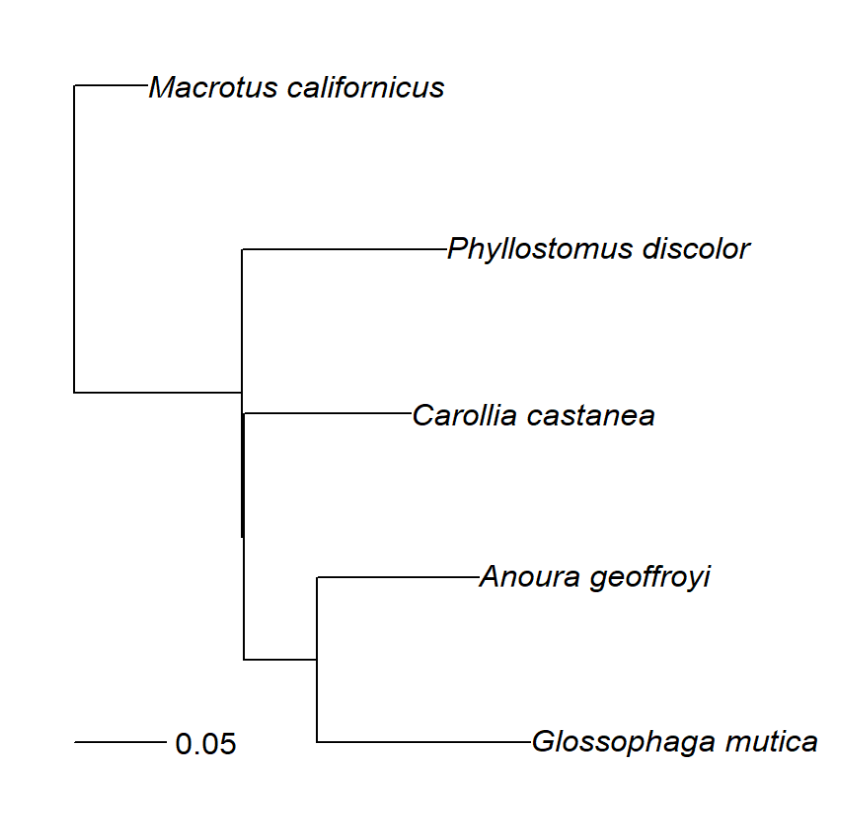

**Figure S1.** Phylogeny based on *cytochrome B* sequences used for phylogenetic statistical corrections. Branch length scale is in number of nucleotide substitutions
