## Supplemental Table Descriptions for "Novel modifications to fructose metabolism are associated with adaptation to high-sugar diets in bats"

Table S1 -  Number of individuals sampled per timepoint for glucose and fructose fed conditions separated by sex
Table S2 -  NCBI accessions used in phylogeny generation
Table S3 -  ANOVA p-values for initial metabolite concentrations
Table S4 -  Pairwise t-test p-values for initial metabolite concentrations per species pooled between feeding groups
Table S5 -  Pairwise phylogenetic t-test p-values for initial metabolite concentrations per species pooled between feeding    	    groups
Table S6 -  ANOVA p-values for species, timepoint, and interaction after glucose feeding
Table S7 -  Pairwise t-test p-values for change in metabolite concentrations between timepoints across species after 	 	    glucose feeding
Table S8 -  Pairwise t-test p-values for change in metabolite concentrations between timepoints within each species 	  	    after glucose feeding
Table S9 -  Pairwise t-test p-values for differences in metabolite concentrations between species across timepoint after 	    glucose feeding
Table S10 - Pairwise phylogenetic t-test p-values for differences in metabolite concentrations between species across 	  	    timepoint after glucose feeding
Table S11 - Pairwise t-test p-values for differences in metabolite concentrations between species at timepoint 0 after 	    	    glucose feeding
Table S12 - Pairwise phylogenetic t-test p-values for differences in metabolite concentrations between species at 	  	    timepoint 0 after glucose feeding
Table S13 - Pairwise t-test p-values for differences in metabolite concentrations between species at timepoint 5 after 	    	    glucose feeding
Table S14 - Pairwise phylogenetic t-test p-values for differences in metabolite concentrations between species at 	  	    timepoint 5 after glucose feeding
Table S15 - Pairwise t-test p-values for differences in metabolite concentrations between species at timepoint 10 after 	    glucose feeding
Table S16 - Pairwise phylogenetic t-test p-values for differences in metabolite concentrations between species at 	  	    timepoint 10 after glucose feeding
Table S17 - ANOVA p-values for sugar, timepoint, and interaction for each species
Table S18 - ANOVA p-values for species, timepoint, and interaction after fructose feeding
Table S19 - Pairwise t-test p-values for change in metabolite concentrations between timepoints across species after 	 	    fructose feeding
Table S20 - Pairwise t-test p-values for change in metabolite concentrations between timepoints within each species 	  	    after fructose feeding
Table S21 - Pairwise t-test p-values for differences in metabolite concentrations between species across timepoint after 	    fructose feeding
Table S22 - Pairwise phylogenetic t-test p-values for differences in metabolite concentrations between species across 	  	    timepoint after fructose feeding
Table S23 - Pairwise t-test p-values for differences in metabolite concentrations between species at timepoint 0 after 	    	    glucose feeding
Table S24 - Pairwise phylogenetic t-test p-values for differences in metabolite concentrations between species at 	  	    timepoint 0 after glucose feeding
Table S25 - Pairwise t-test p-values for differences in metabolite concentrations between species at timepoint 5 after 	    	    glucose feeding
Table S26 - Pairwise phylogenetic t-test p-values for differences in metabolite concentrations between species at 	  	    timepoint 5 after glucose feeding
Table S27 - Pairwise t-test p-values for differences in metabolite concentrations between species at timepoint 10 after 	    glucose feeding
Table S28 - Pairwise phylogenetic t-test p-values for differences in metabolite concentrations between species at 	  	    timepoint 10 after glucose feeding
